# Modulation of proteasome activity by curcumin and didemethylcurcumin

**DOI:** 10.1101/2020.07.27.222679

**Authors:** Tapan K. Khan, Youngki You, Thomas J. Nelson, Subrata Kundu, Saroj Pramanik, Joydip Das

**Affiliations:** Center for Neurodegenerative Diseases, Blanchette Rockefeller Neurosciences Institute, West Virginia University, Morgantown, WV 26506, USA; Department of Pharmacological and Pharmaceutical Sciences, College of Pharmacy, University of Houston, Houston, TX 77204; Department of Chemistry, Indian Institute of Technology Delhi, New Delhi, Delhi 110016, India; Department of Biology, Morgan State University, Baltimore, MD 21251; Department of Veterans Affairs, Rocky Mountain Mental Illness, Research, Education and Clinical Care, Denver, Aurora, CO 80045; Department of Neurology, Marshall University School of Medicine, Huntington WV 25704

**Keywords:** Curcumin, Didemethylcurcumin, Proteasome, Molecular Docking, Neurodegeneration, Cancer

## Abstract

Modulation of proteasome function by pharmacological interventions and molecular biology tools is an active area of research in cancer biology and neurodegenerative diseases. Curcumin (diferuloylmethane) is a naturally occurring polyphenol that affects multiple signaling pathways and known to modulate PKC activities. The therapeutic significance of curcumin is often considered to reside in its anti-inflammatory, antioxidant, anti-angiogenic, or anti-apoptotic properties. However, recent research suggests that the therapeutic efficacy of curcumin may be due to its activity as a potent inhibitor of the proteasome. In this study, we show that both curcumin and its synthetic polyphenolic derivative (didemethylcurcumin), CUIII modulated proteasome activity in a biphasic manner. Curcumin and CUIII increased proteasome activity at nanomolar concentrations but inhibited proteasome activity at micromolar concentrations. Curcumin was more effective than CUIII in relative proteasome activity increase at nanomolar concentrations. Also, curcumin was more effective than CUIII in relative proteasome activity inhibition at micromolar concentration. The docking study was conducted on the 20S proteasome catalytic subunit. Estimated Kd values for curcumin and didemethylcurcumin are 0.0054μM and 1.3167μM, respectively. These values correlate well with the results of the effectiveness of modulation by curcumin compare to CUIII. The small size of CUIII makes its dock to the narrow cavity of the active site, but the binding interaction is not strong compare to curcumin. This study suggests that curcumin and its didemethyl derivative can be used to modulate proteasome activity. This communication implicates the reason why curcumin and its didemethyl derivative can be used to two different disease mechanisms, neurodegeneration, and cancer.

## Introduction

Proteasome degrades and clears aggregated, damaged, and misfolded proteins in ubiquitin-proteasome pathways that control cellular homeostasis and proteostasis (Goldberg, 2003; Hershko and Ciechanover, 1998). Aging, age-related neurodegenerative diseases, and cellular senescence decrease proteasome activity. In contrast, proteasome activity is increased in cancer cells. Proteasome activity inhibitors exert anti-apoptotic properties that can prevent angiogenesis and metastasis. Proteasome activity is reduced in neurodegenerative diseases, including Alzheimer’s disease, Parkinson’s disease, Huntington’s disease, and amyotrophic lateral sclerosis (Jansen et al., 2014; Riederer et al., 2011; Saez, and Vilchez, 2014; Lin, and Tan, 2007; Thibaudeau et al., 2018). The development of proteasome activators for the potential therapeutic agents to treat neurodegenerative diseases is based on the principle that increased proteasome activity can increase the rate of degradation of different neurotoxic aggregated, misfolded, oxidized buildup in brain cells. Proteasome inhibitors have a long history in cancer therapy. Reduced proteasome activity can reduce the survival of malignant cells by inducing apoptosis and inhibit NF-kappa B transcriptional factor that inhibitors can prevent angiogenesis and cellular invation in vivo (Almond and Cohen, 2002). Patients with myeloma showed initial positive responses by the treatment with proteasome inhibitors (Manasanch and Orlowski, 2017).

Natural polyphenols and its simple analogs with improved bioavailability have the potential to be drugs for cancer and neurodegenerative diseases in the future. Several polyphenols exert their antioxidant properties by regulating the gene expression and signaling pathways. The two most important polyphenols, curcumin, and resveratrol are implicated in therapeutics of diseases like cancer and Alzheimer’s disease. Curcumin has been tested in diagnosis, prevention, and treatment of Alzheimer’s disease (Ringman et al., 2012; Koronyo et al., 2017; den Haan et al., 2018; Chen et al., 2018). It has also been tested in clinical trials for cancer, Alzheimer’s disease, and inflammatory diseases.

The objective of the present study is to understand if curcumin and its derivatives exert their therapeutic actions by modulating activity of proteasome. This study reports how the proteasome function can be modulated by curcumin and its synthetic polyphenolic derivative. At lower concentrations (nanomolar), both curcumin and its synthetic polyphenolic derivative increased proteasome activity. By contrast, at the micromolar level, both decreased proteasome activity. Therefore, both curcumin and its synthetic polyphenolic derivative can be used as a therapy in neurodegenerative conditions. On the other hand, both compounds may be effective for cancer therapy at micromolar concentrations.

## Experiments

### Materials and Methods

#### Synthesis of didemethylcurcumin (CUIII) from Curcumin (CU)

The synthesis of polyphenolic curcumin derivative, tetrahydroxy curcumin ((1E, 4Z, 6E)-1,7-bis(3, 4-dihydroxyphenyl)-5-hydroxyhepta-1, 4, 6-trien-3-one; CUIII) is described in the literature (Majhi et al., 2010). Briefly, the synthesis of the CUIII was initiated from curcumin, which was obtained by the recrystallization of commercially available (purity75–80%) curcumin (Sigma-Aldrich). Bromination reaction was performed by boron tribromide in anhydrous dichloromethane at −78°C to remove two methoxy groups. In the next step, bromide groups were replaced by hydroxyl groups using saturated sodium bicarbonate solution with stirring. CUIII was purified from the reaction mixture by column chromatography (Hexane: EtOAc: MeOH; 60:38:2 %). The molecular structure of CUIII was authenticated by NMR and mass spectral analysis. The reaction scheme and molecular structure of CUIII are depicted in Figure 1. The stock solution of differentiating reagent, retinoic acid (Sigma-Aldrich) was prepared in ethanol and kept in dark at 4°C until used in cell culture.

**Figure 1:**
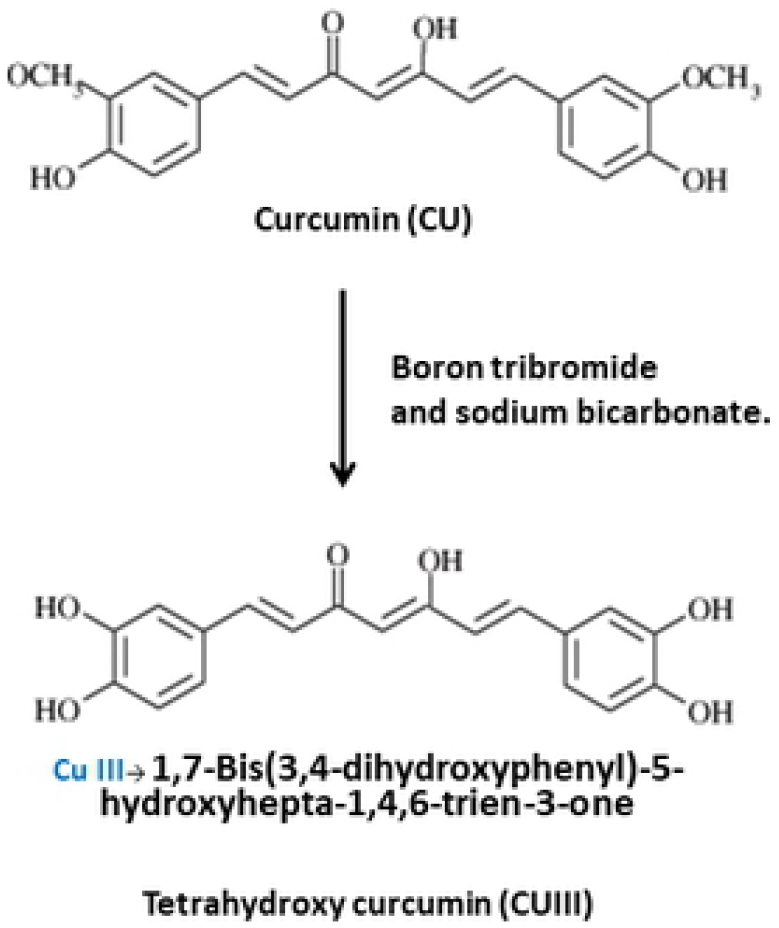
Synthetic scheme for polyphenolic analog, didemethylcurcumin (CUIII) from curcumin (CU).

#### Cell culture

Commercially available Human SH-SY5Y neuroblastoma cell line was obtained from Sigma-Aldrich. Cells were cultured and maintained in culture medium containing 45% MEM (Minimum Essential Medium Eagle), 45% F-12K (Nutrient Mixture F-12 Ham, Sigma-Aldrich) and 10% FBS (Atlanta Biologicals). The tissue culture incubator was monitored at 37°C, 5% CO2, and 90% humidity. Retinoic acid solution (final concentration 10 μM in culture medium) was added directly into ~80-90% confluence cells. After two days, cells were differentiated into neuron-like cells. We called transformed SH-SY5Y cells. Those transformed cells were recultured in reduced serum medium (49% F-12K, 49% MEM and 2% FBS) until ready for treatment.

#### Preparation of cell lysate and treatments with Curcumin (CU) and didemethylcurcumin (CUIII)

Stock solutions of curcumin (CU) and didemethylcurcumin (CUIII) were prepared in ethanol and stored in the dark at −20°C. Differentiated neuron-like SH-SY5Y cells were treated with different concentrations (0, 20 nM, and 20 μM) of CU and CUIII for 24 hours. Transformed SH-SY5Y cells were harvested by washing three times with 1×PBS (4°C, pH=7.4) and homogenized by a cell homogenizer in 1×PBS (4°C, pH=7.4) containing 0.5% NP-40. Homogenized cells were sonicated by sonication at 4°C. After sonication, supernatants were collected by centrifugation and stored as cell lysate. Proteasome activity assay was conducted on each supernatant. Duration of sonication had no effect on proteasome activity assay. Protein concentrations in each experiment were determined routinely using Bradford reagent (Cat# 23238; Thermo Scientific, Rockford, IL) by following manufacturer protocol.

### Measurement of proteasome activity using fluorometric assay

Cell lysate was added in 0.5 mL of proteasome assay buffer (Cat#K245-100-1; supplied by BioVision, Milpitus, CA) in the fluorescence microcuvette. The calculated amount of cell lysate was added to make a final protein concentration of 0.5mg/mL in the fluorescence microcuvette. The actual procedure of measurement of proteasome activity using fluorometric assay has been published elsewhere (Khan and Nelson, 2018). In principle, the assay monitored the chymotrypsin-like activity of the 20S proteasome assembly using a 7-amino-4-methylcoumarin (AMC)-tagged peptide substrate (Succ-LLVY-AMC) (BioVision (Milpitus, CA). Highly fluorescent AMC molecules were released from the reaction in the presence of proteolytic cleavage by the existing proteasome in the cell lysate remained in the fluorescence cuvette. Fluorogenic substrate, AMC-tagged peptide has been utilized for 20S proteasome activity assay extensively in literature. Fluorescence intensity of AMC was measured as a function of time and monitored by a fluorescence spectrometer (Photon Technology International, Inc. Birmingham, NJ). Fluorometric intensity data were collected as a function of time using a 2-nm bandwidth. Highly stable illumination with very low ripple is required to reduce noise in fluorescence intensity measurements. We modified the light source of the Photon Technology International fluorescence spectrometer. An 75W xenon arc lamp powered by an OPS-A150 power supply was used as a light source (Newport Corp., Irvine, CA). All reagents related fluorometric measurements, such as assay buffer, AMC-tagged peptide substrate (Succ-LLVY-AMC), proteasome activity inhibitor (Carbobenzoxy-Leu-Leu-leucinal; MG-132), and Jurkat Cell lysate were purchased from BioVision (Milpitas, CA). In actual measurement, the time dependent proteolytic activity was monitored by the fluorescent AMC production at Ex/Em=350/440 nm in a temperature-controlled fluorescence cuvette setup. The whole proteolytic activity measurement was monitored for one hour data acquisition. Protocol of proteasome activity fluorometric assay kit (Ct# K245-100; BioVision, CA) suggested that initial AMC fluorescence intensity reading (~20-30 min) would not be linear. Therefore, the linear regression was conducted on data collected in the last 30 min in data analysis. The slope of the kinetic was estimated by the linear regression of data collected within 30 to 60 min and analyzed by Sigma Stat software (Systat Software, Inc., San Jose, CA).

The procedure of standardization of the proteasome activity assay has been published elsewhere (Khan and Nelson, 2018). Briefly, standardizing reagent, we used a positive control sample (Jurkat Cell lysate; BioVision, Milpitas, CA) with significant proteasome activity for standardization of the fluorescence spectrometer. The standard proteasome inhibitor, Carbobenzoxy-Leu-Leu-leucinal (MG-132; BioVision, Milpitas, CA), was used to reduce the proteasome activity. MG-132 has been used routinely as a potential proteasome inhibitor in the literature (Pastore et al., 2013; Orthwein et al., 2015; Ma et al., 2015). We checked the specificity of fluorescence intensity accumulation due to proteasome activity by adding MG-132 in fluorescence measuring cuvette. This experiment determined whether fluorescence intensity accumulating in reaction mixture directly originating from proteasome activity only, not from other protease activities.

#### Molecular Docking

Curcumin and didemethylcurcumin were docked into 20S proteasome using Sybyl X 2.1 (Certara Inc., Princeton, NJ). X-ray crystal structure of human 20S proteasome (PDB: 5LE5) was used for the protein structure, and the curcumin and didemethylcurcumin were downloaded from PubChem. The energy of 20S proteasome applied by AMBER7 FF99 force field was minimized using Powel method. Tripos force field was applied to the two ligands, and then they were stabilized using Powel energy minimization method. For the molecular docking, protomols were generated by Sybyl for a docking space of ligand by selecting the specific residues within 0.1 Å of a known 20S proteasome inhibitor, bortezomib (PDB: 5LF3) (Schrader et al., 2016). Threshold 0.2 and Bloat 1.0 were used to create the protomols. After the protomol had been generated, ligand docking was performed using the Surflexdock Geom module of Sybyl. For the analysis, structures were visualized using Discovery Studio Visualizer 4.5 (Biovia Inc., San Diego, CA).

#### Data analysis

Systat software (Systat Software, Inc., San Jose, CA), and Microsoft Excel were used for all statistical analyses. Proteasome activity for each experiment was determined by performing linear regression analysis on fluorescence intensity data collected in the time range of 30-60 min. P values were estimated by Student T-test. T-test type: paired, two-sample equal variance.

## Results and discussion

### Modulation of proteasome activity by curcumin (CU), and didemethylcurcumin (CUIII) treatment

We examined the effect of CU and the CUIII treatment on 20S proteasome catalytic subunit activity in differentiated cultured neuron-like SH-SY5Y cells (T-SH-SY5Y). We tested 20S proteasome activity at two different concentrations of CU and CUIII: 20nM and 20μM. T-SHSY-5Y cells were cultured and treated with vehicle (Control), and two widely different concentrations of CU, and CUIII for 24 hours. Proteasome activities in cell lysates were monitored by measuring the accumulation of fluorescence intensity (A.U.) of highly fluorescent AMC (7-amino-4-methylcoumarin=AMC) molecules due to proteolytic cleavage of the substrate Succ-LLVY-AMC in the fluorescence micro-cuvette (Figure 2A). The time-base fluorescence intensity measurement setup was fixed at Ex/Em=350/440 nm. Relative proteasome activities were calculated in cell lysates prepared from T-SHSY-5Y cells treated with vehicle (Control), CU, and CUIII. (Figure 2B) normalizing with Control as 100%. We examined whether resulting fluorescence intensity accumulation in the fluorescence microcuvette was due to the proteolytic cleavage of the substrate Succ-LLVY-AMC by proteasome activity. When we added proteasome inhibitor MG-132 in the fluorescence microcuvette in each experiment and found MG-132 completely suppressed the increased fluorescence intensity (data not are shown), those experiments suggested that resulting fluorescence intensity increase was only due to the proteasome activity. Here we actually measured the chymotrypsin-like activity of 20S proteasome subunit in the presence of CU and CUIII treatment with different concentrations added in neuron like T-SH-SY5Y cells. The relative proteasome activity increased by nanomolar (nM) concentration and decreased at micro-molar (20 μM) concentration treatment. The relative proteasome activity was calculated from the time dependent fluorescence increase with a normalization with the vehicle (Control) as 100%. The relative proteasome activity in the presence of 20μM CU and CUIII was lower than the vehicle control. In contrast, the relative proteasome activity at 20nM CU and CUIII were significantly higher than that of control (Figure 2). We found no specific proteasome activity after the addition of proteasome inhibitor MG-132. CU and CUIII both increased proteasome activity at nanomole concentration, but inhibited proteasome activity at 20 μM concentration (Figure 2B). Error bars were calculated as the mean of standard errors from three independent experiments. CU was more effective than CUIII in relative proteasome activity increase by nanomolar concentration (CU vs CUIII p< 0.003). Also, CU was more effective than CUIII in relative proteasome activity inhibition by micromolar concentration (CU vs CUIII p< 0.0003).

**Figure 2:**
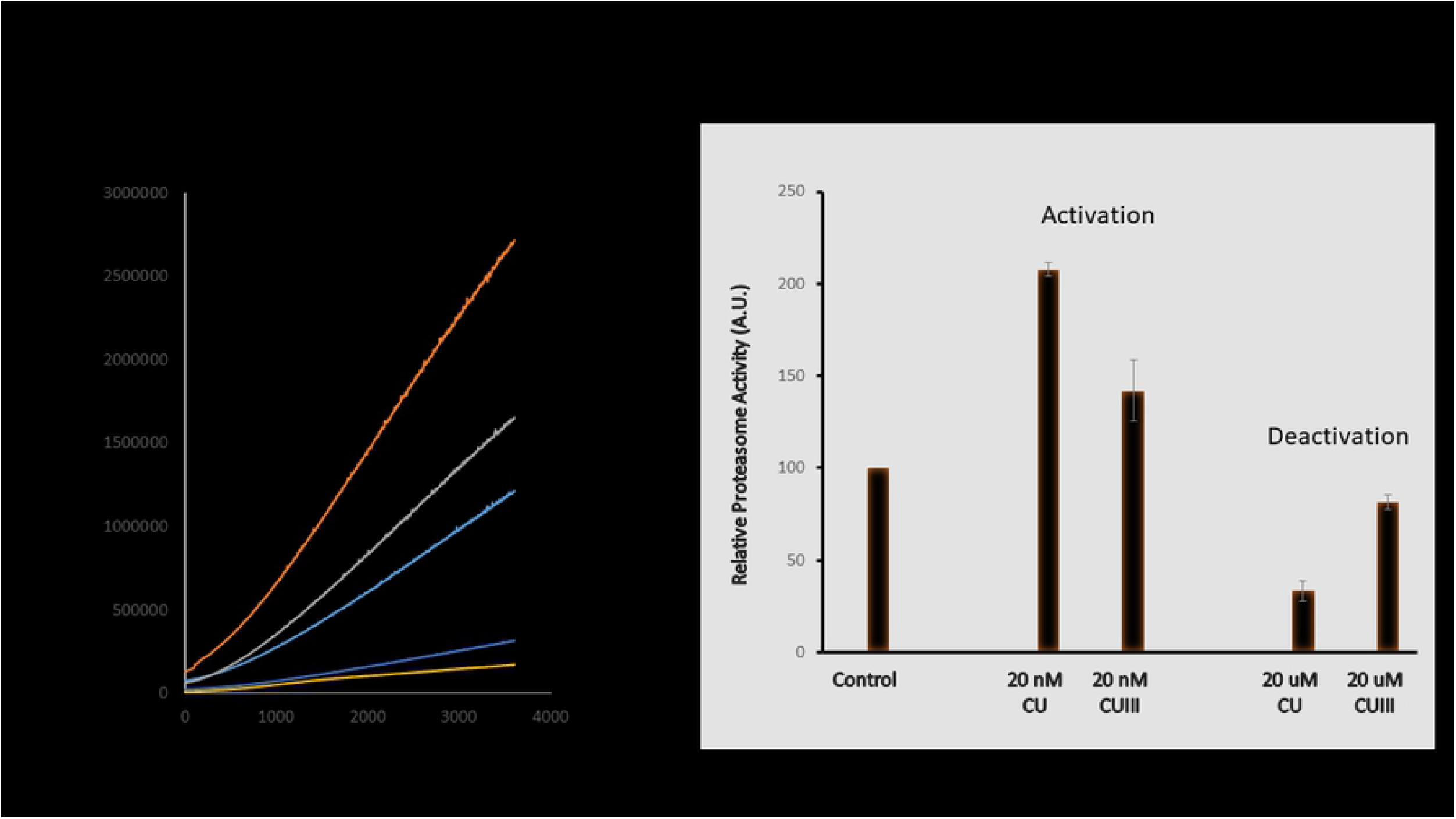
Curcumin (CU) and its polyphenolic derivative, didemethylcurcumin (CUIII) modulate proteasome activity in differentiated neuron-like SH-SY5Y cells (T-SHSY5Y). Fluorescence intensity accumulation as a function of time due to AMC released by the proteasome activity in a fluorescence micro-cuvette containing cell lysates from T-SHSY5Y cells. Those cells are previously cultured in presence of vehicle (Control), and different concentrations of curcumin (CU) and didemethylcurcumin (CUIII) for 24 hours (A). Proteasome activity in the presence of vehicle (Control), 20 nM curcumin (CU), 20 nM didemethylcurcumin (CUIII), and 20 μM curcumin (CU), 20 μM didemethylcurcumin (CUIII) treatment (B). Relative proteasome activity increases by nano molar (nM) concentration and decreases at micro-molar (20 μM) concentration treatment. Error bar =standard error of means from three independent measurements. 7-amino-4-methyl coumarin=AMC.

### Docking of curcumin (CU) and didemethylcurcumin (CUIII) to proteasome

To understand the binding site of curcumin and didemethylcurcumin and the interactions between the human 20S proteasome and the ligands, curcumin and didemethylcurcumin were docked into β-5 subunit and β-6 subunit of 20S proteasome. The two compounds were docked into the chymotrypsin active site of proteasome as curcumin showed the inhibitory effect on the chymotrypsin activity of a purified rabbit proteasome and human 26S proteasome. The parameters for molecular docking were verified by docking the known inhibitor, bortezomib, into the active site. The docking pose of bortezomib was similar to the binding pose of bortezomib in the cocrystal structure (Figure 3A). Both curcumin and didemethylcurcumin were also docked into the active site like bortezomib (Figure 3B and C). The hydroxyl groups on a phenol ring of two compounds interacted with Thr-1 of β-5 subunit. However, the hydroxyl groups on the other side of phenol ring of curcumin and didemethylcurcumin bound to different residues, Ala-50 of β-5 subunit and Gln-131 of β-6 subunit, respectively (Figure 4). Didemethylcurcumin docked into the small pocket surrounded by β-5 subunit and β-6 subunit together (Figure 3C). But, it was not shown in the curcumin because of the steric conflicts between the methoxyl groups on phenol rings and the residues, Arg-19, Tyr130, and Gln-131 (Figure 3B). Although curcumin did not bind to the small pocket at β-5 and β-6 subunit, it showed a higher affinity to the proteasome than didemethylcurcumin. The estimated Kd values from the docking score of curcumin and didemethylcurcumin are 0.0054 μM (docking score, 8.2705) and 1.3167 μM (docking score, 5.8805), respectively (Jain, 1996). Indeed, curcumin formed more number of hydrogen bonds with the proteasome than didemethylcurcumin (Figure 4). The hydroxyl and methoxyl groups of curcumin formed six hydrogen bonds with Thr-1 (amine and hydroxyl group), Thr-21, Ala-49, Als-50, and Asp-125. In didemethylcurcumin, the hydroxyl groups formed four hydrogen bonds with Thr-1, Arg-19, Gln-131, and Tyr-169. Under similar docking conditions, a Kd value of 0.0537 μM (docking score, 7.2702) was obtained for bortezomib that forms a covalent bond with the Thr1 residue of proteasome.

**Figure 3.**
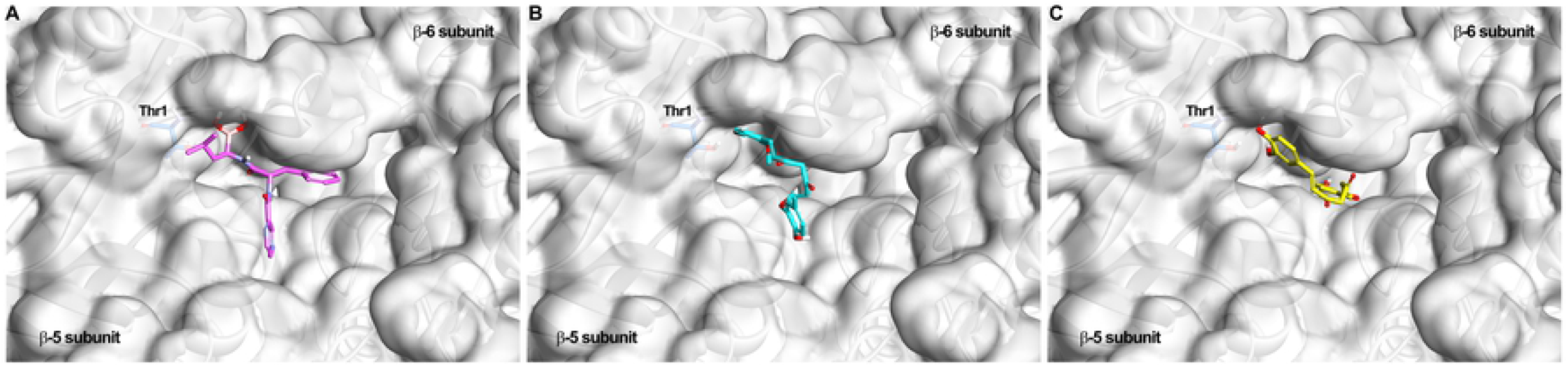
Docking of curcumin, didemethylcurcumin, and bortezomib to the 20S proteasome. Top-scoring binding poses of (A) bortezomib, (B) curcumin, and (C) didemethylcurcumin. Curcumin and didemethylcurcumin were docked to proteasome like bortezomib. However, a hydroxyl group on phenol ring of curcumin and didemethylcurcumin bound to different sites. Thr-1 is represented by blue colored stick. The surface of β-5 subunit and β-6 subunit are colored by gray. β-5 subunit and β-6 subunit are represented by the solid ribbon.

**Figure 4.**
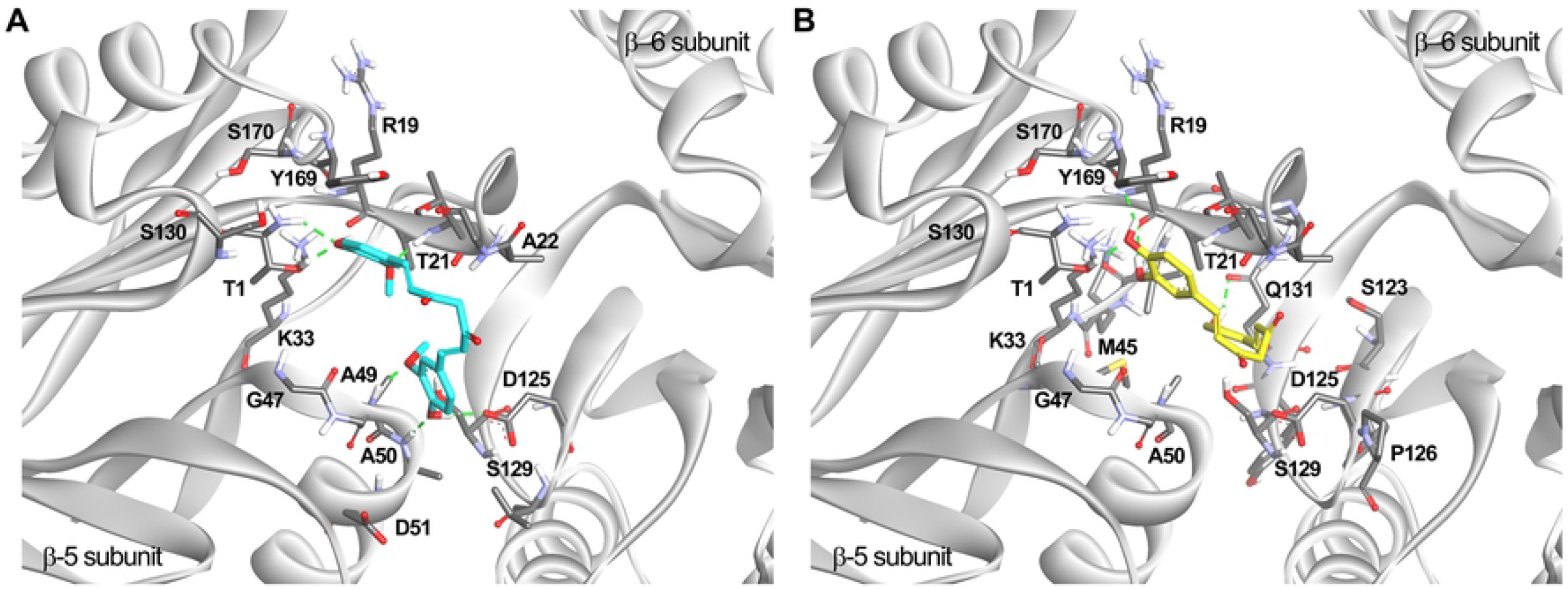
Docking of curcumin and didemethylcurcumin to the 20S proteasome. Top-scoring binding poses of (A) curcumin and (B) didemethylcurcumin. The hydroxyl and methoxyl groups of curcumin formed six hydrogen bonds and didemethylcurcumin formed four hydrogen bonds. Only residues within 5 Å of the compounds are reported. Hydrogen bonds are shown as dotted lines (in green). β-5 subunit and β-6 subunit are represented by the solid ribbon (in gray).

In summary, both curcumin and didemethylcurcumin bound to the Thr-1 of chymotrypsin active site of proteasome. However, didemethylcurcumin reached more toward β-6 subunit and formed less number of hydrogen bonds than curcumin. As a result, curcumin showed higher affinity to the proteasome than didemethylcurcumin.

In literature, the therapeutic significance of curcumin is often considered to reside in its anti-inflammatory, antioxidant, anti-angiogenic, or anti-apoptotic properties. However, this study suggests that the therapeutic efficacy of curcumin may be due to its activity as a potent inhibitor of the proteasome. Modulation of proteasome function is implicated in cancer therapy and in reducing neurotoxic aggregated oligomers buildups in the brain in neurodegenerative diseases. Mechanistic analyses suggested that these aggregated oligomers inhibit the 20S proteasome activity through allosteric impairment of the substrate gate in the 20S core particle (Thibaudeau et al., 2018).

Most of the studies found inhibitory effects of curcumin on proteasome activity at higher concentration range. The biphasic effect of curcumin on proteasome activity has been reported in only one study (Ali and Rattan, 2006). Stimulatory effects were shown at low concentrations (nM to μM) and inhibitory effects were observed at >1μM concentration. The effect of curcumin and its poly phenolic derivative (CUIII) on proteasome activity appears to be dose-dependent, as low doses (nM) increase proteasome activity whereas high doses (μM) inhibit the proteasome activity. The inhibition of proteasome at high concentration is well known. Moreover, for neurodegenerative disease, it requires increasing proteasome activity to reduce unwanted protein aggregates.

In our molecular docking studies, the docking pose of proteasome inhibitor, bortezomib was not same with the conformation of bortezomib in the cocrystal structure because it binds to Thr-1 irreversibly in the crystal structure (Figure. 3). However, the overall conformation of docked bortezomib was similar to the bortezomib in the crystal structure.

Milacic et al. reported molecular docking of curcumin using Autodock (Milacic et al., 2008). They focused on the carbonyl carbon which conferred proteasome-inhibitory potencies (Milacic et al., 2008; Daniel et al., 2006). In their study, the carbonyl carbon of curcumin bound to nearby Thr-1 and a hydroxyl group of curcumin bound to Ser-96 of β-5 subunit. The binding site is not similar to the active site of bortezomib. There was limited resource for protein structure of proteasome. The protein structure they used was from yeast, and the resolution was 3.0 Å (Smith et al., 2004; Groll et al., 2001). It might be the reason why the docking pose of curcumin in our study was different from the docking pose in their study.

The therapeutic significance of concentration dependence modulation of proteasome activity by curcumin has not been implicated concerning different disease mechanisms, such as Alzheimer’s disease and cancer. Concentration-dependent modulation of proteasome function has therapeutic significance in neurodegenerative diseases, including Alzheimer’s disease. This study implicates that both curcumin and its polyphenolic derivative can be used to activate proteasome function at nanomolar concentration. Toxic aggregated and misfolded proteins in neurodegenerative diseases can be removed from the brain cells at lower concentrations (nanomolar). Therefore, at lower concentration curcumin and its polyphenolic derivative may be used for neuroprotection studies in future studies in different animal models of neurodegenerative diseases. In contrast, at micromolar concentration curcumin and its polyphenolic derivative can decrease proteasome activity.

Curcumin has very low bioavailability (Anand et al., 2008). Pharmacokinetic results of a clinical trial of Alzheimer’s disease found very low levels of curcumin in blood plasma at the 24-week visit (2.67±1.69 ng/mL of curcumin and 2.67±1.69 ng/mL of its metabolic product, tetrahydrocurcumin) (Ringman et al., 2012). The study also found no detectable levels of curcumin in cerebrospinal fluid. Curcumin and CUIII need to have sufficient bioavailability to reach the concentrations necessary to inhibit the proteasome, especially if they require micromolar concentrations to work. But structural analogs and liposomal formulations have greatly improved it.

## Author contributions

TKK and JD conceived and supervised the study; TKK and JD designed experiments; TKK and YY performed experiments; TJN, SP, and SK provided new tools and reagents; TKK and JD analyzed data; TKK, JD and YY wrote the manuscript; TKK and JD made manuscript revisions.

## Conflict of Interest

Authors declare no conflicts of interest and their respective Institutes have no other agreements and financial interest in this work.

## Acknowledgments

The research was partly supported by the Intramural Research Program of The Rockefeller Neuroscience Institute (formerly, Blanchette Rockefeller Neurosciences Institute) at West Virginia University, Morgantown, WV (T.K.K.) and funding from National Institutes of Health Grant 1R01 AA022414 to J.D.

## Reference

Ali RE, Rattan SI. Curcumin’s biphasic hormetic response on proteasome activity and heat-shock protein synthesis in human keratinocytes. Annals of the New York Academy of Sciences, 2006; 1067: 394–399. DOI: 10.1196/annals.1354.056.

Almond JB, Cohen GM. The proteasome: a novel target for cancer chemotherapy. Leukemia, 2002;16:433–443.

Anand P, Thomas SG, Kunnumakkara AB, Sundaram C, Harikumar KB, Sung B, Tharakan ST, Misra K, Priyadarsini IK, Rajasekharan KN, Aggarwal BB. Biological activities of curcumin and its analogues (Congeners) made by man and Mother Nature. Biochemical Pharmacology, 2008; 76:1590–1611.

Chen M, Du Z-Y, Zheng X, Li D-L, Zhou R-P, Zhang K. Use of curcumin in diagnosis, prevention, and treatment of Alzheimer’s disease. Neural Regen Res, 2018; 13(4): 742–752. doi: 10.4103/1673-5374.230303.

Daniel KG, Landis-Piwowar KR, Chen D, Wan SB, Chan T-H, Dou QP. Methylation of green tea polyphenols affects their binding to and inhibitory poses of the proteasome β5 subunit. International Journal of Molecular Medicine, 2006; 18: 625–632.

den Haan J, Morrema THJ, Rozemuller AJ. et al. Different curcumin forms selectively bind fibrillar amyloid beta in post mortem Alzheimer’s disease brains: Implications for in-vivo diagnostics. Acta Neuropathol commun, 2018; 6: 75. https://doi.org/10.1186/s40478-018-0577-2.

Goldberg AL. Protein degradation and protection against misfolded or damaged proteins. Nature, 2003; 426: 895–899.

Groll M, Koguchi Y, Huber R, Kohno J. Crystal structure of the 20 S proteasome: TMC-95A complex: a non-covalent proteasome inhibitor. Journal of Molecular Biology, 2001; 311: 543–548.

Hershko A, Ciechanover A. The ubiquitin system. Ann Rev Biochem, 1998; 67: 425–479.

Jain AN. Scoring noncovalent protein-ligand interactions: a continuous differentiable function tuned to compute binding affinities. Journal of computer-aided molecular design, 1996; 10: 427–440.

Jansen AH, Reits EA, Hol EM. The ubiquitin proteasome system in glia and its role in neurodegenerative diseases. Frontiers Molecular Neuroscim 2014; 7:73. doi: 10.3389/fnmol.2014.00073.

Khan TK, Nelson TJ. Protein kinase C activator bryostatin-1 modulates proteasome function. J Cell Biochem, 2018;119(8):6894–6904. doi: 10.1002/jcb.26887.

Koronyo Y, Biggs D, Barron E, Boyer DS, Pearlman JA, Au WJ, Kile SJ, Blanco A, Fuchs DT, Ashfaq A, Frautschy S, Cole GM, Miller CA, Hinton DR, Verdooner SR, Black KL, Koronyo-Hamaoui M. Retinal amyloid pathology and proof-of-concept imaging trial in Alzheimer’s disease. JCI Insight, 2017; 2(16): e93621. doi: 10.1172/jci.insight.93621.

Lim K-L, Tan JMM. Role of the ubiquitin proteasome system in Parkinson’s disease. BMC Biochem, 2007; 8(Suppl 1): S13. doi: 10.1186/1471-2091-8-S1-S13.

Ma B, Chen Y, Chen L, Cheng H, Mu C, Li J, Gao R, Zhou C, Cao L, Liu J, Zhu Y, Chen Q, Wu S. Hypoxia regulates Hippo signaling through the SIAH2 ubiquitin E3 ligase. Nat Cell Biol. 2015; 17: 95–103.

Majhi A, Rahman GM, Panchal S, Das J. Binding of curcumin and its long chain derivatives to the activator binding domain of novel protein kinase C. Bioorg Med Chem, 2010; 18(4):1591–8. doi: 10.1016/j.bmc.2009.12.075.

Manasanch EE, Orlowski RZ. Proteasome inhibitors in cancer therapy. Nature Rev Clinic Oncol, 2017;14:417–433.

Milacic V, Banerjee S, Landis-Piwowar KR, Sarkar FH, Majumdar AP, Dou QP. Curcumin inhibits the proteasome activity in human colon cancer cells in vitro and in vivo. Cancer research, 2008; 68: 7283–7292.

Orthwein A, Noordermeer SM, Wilson MD, Landry S, Enchev RI, Sherker A, Munro M, Pinder J, Salsman J, Dellaire G, Xia B, Peter M, Durocher D. A mechanism for the suppression of homologous recombination in G1 cells. Nature, 2015; 528:422–426.

Pastore N, Blomenkamp K, Annunziata F, Piccolo P, Mithbaokar P, Maria Sepe R, Vetrini F, Palmer D, Ng P, Polishchuk E, Iacobacci S, Polishchuk R, Teckman J, Ballabio A, Brunetti-Pierri N. Gene transfer of master autophagy regulator TFEB results in clearance of toxic protein and correction of hepatic disease in alpha-1-anti-trypsin deficiency. EMBO Mol Med, 2013; 5:397–412.

Riederer BM, Leuba G, Vernay A, Riederer IM. The role of the ubiquitin proteasome system in Alzheimer’s disease. Exp Biology Med (Maywood), 2011; 236:268–276.

Ringman JM, Frautschy SA, Teng E, Begum AN, Bardens J, Beigi M, Gylys KH, Badmaev V, Heath DD, Apostolova LG, Porter V, Vanek Z, Marshall GA, Hellemann G, Sugar C, Masterman DL, Montine TJ, Cummings JL, Cole GM. Oral curcumin for Alzheimer’s disease: tolerability and efficacy in a 24-week randomized, double blind, placebo-controlled study. Alzheimer’s Research & Therapy, 2012; 4: 43. https://doi.org/10.1186/alzrt146.

Saez I, Vilchez D. The mechanistic Links between proteasome activity, aging and age-related Diseases. Current Genomics, 2014; 15:38–51.

Schrader J, Henneberg F, Mata RA, Tittmann K, Schneider TR, Stark H, Bourenkov G, Ashwin C. The inhibition mechanism of human 20S proteasomes enables next-generation inhibitor design. Science 2016; 353: 594–598.

Smith DM, Daniel KG, Wang Z, Guida WC, Chan T-H, Dou QP. Docking studies and model development of tea polyphenol proteasome inhibitors: applications to rational drug design. Proteins: Structure, Function, and Bioinformatics, 2004; 54: 58–70.

Thibaudeau TA, Anderson RT, Smith DM. A common mechanism of proteasome impairment by neurodegenerative disease-associated oligomers. Nature Commun, 2018; 9:1097.

